# B7-H4 enrichment represents a site-specific immunotherapy target in small bowel gastrointestinal stromal tumors

**DOI:** 10.64898/2026.08.04.742601

**Authors:** Henry Singer, Montana Morris, Roberta Maestro, Angelo Paolo Dei Tos, Ronald P. DeMatteo, Gerardo A. Vitiello

**Author notes:** **Corresponding author:** Gerardo Vitiello, 350 Community Drive, Manhasset, NY 11030.

## Abstract

Small bowel gastrointestinal stromal tumors (GISTs) are more aggressive than gastric GISTs, yet the biologic basis for this difference remains poorly understood. We hypothesized that differential expression of immune checkpoints contributes to this site-specific behavior. Bulk RNA sequencing of 42 primary GISTs (36 gastric, 6 small bowel) revealed marked upregulation of *VTCN1*, which encodes the inhibitory checkpoint B7-H4, in small bowel tumors (log_2_FC = 7.95, adjusted *P* < 0.001). In contrast, expression of the therapeutically targeted checkpoints PD-L1, PD-1, and CTLA-4 was comparable between sites. Concordantly, B7-H4 enrichment was accompanied by an immunosuppressive tumor microenvironment, characterized by reduced antigen-presenting cells, fewer effector-memory CD8^+^ T cells, lower granzyme B expression, and suppression of interferon and inflammatory signaling pathways. Notably, the differences in B7-H4 expression were independent of imatinib-treatment status. These findings were corroborated in an external cohort of 77 untreated GISTs, in which *VTCN1* was similarly enriched in small bowel tumors. Independent immunohistochemical analysis of a tissue microarray comprising 68 untreated primary GISTs confirmed the pattern, showing median B7-H4 positivity of 78.6% in duodenal, 20.5% in jejunal/ileal, and 0% in gastric tumors, with staining localized to tumor cells rather than stroma. Collectively, these data identify B7-H4 as a site-specific feature of small bowel GISTs and a potential therapeutic target for tumors that have not responded to conventional checkpoint blockade.

---

Small bowel gastrointestinal stromal tumors (GISTs) behave more aggressively than gastric GISTs—a site-specific risk reflected in GIST staging alongside tumor size, mitotic rate, and location (1)—yet the biological basis for this aggressiveness is poorly understood. We hypothesized that differential immune checkpoint expression may underlie small bowel GIST aggressiveness. Using our published bulk RNA sequencing data (2), we compared immune-related markers of primary gastric (n=36) and small bowel (n=6) GISTs (**Supplemental Table 1)**. *VTCN1*, which encodes the inhibitory checkpoint protein B7-H4, was upregulated in small bowel GISTs relative to gastric tumors (DESeq2: log_2_FC = 7.95, adjusted *P* < 0.001; **Figure 1A**). Therapeutically targeted checkpoint proteins PD-L1 (*CD274*), PD-1 (*PDCD1*), and CTLA-4 (*CTLA4*) were not upregulated (log_2_FC = −1.82, −1.03, −0.38; adjusted *P* = 0.03, 0.39, 0.77), identifying B7-H4 as a site-specific target. xCell deconvolution showed fewer antigen-presenting cells in small bowel GISTs, including activated dendritic cells (aDC: log_2_FC = −0.76, *P* = 0.030) and M1 macrophages (log_2_FC = −1.56, *P* = 0.028), as well as natural killer T (NKT) cells (log_2_FC = −1.32, *P* = 0.027) (**Figure 1B**). While total CD8+ T-cells were expanded (log_2_FC = 0.40, *P* = 0.035), the CD8 effector memory subset was depleted (CD8+ Tem: log_2_FC = −1.64, *P* = 0.020). Across primary GISTs (n=42), B7-H4 expression was negatively correlated with predicted effector memory populations (CD4+ Tem: ρ = −0.447, *P* = 0.003; CD8+ Tem: ρ = −0.427, *P* = 0.005) and with granzyme B expression (ρ = −0.308, *P* = 0.047), consistent with B7-H4 mediated cell cycle arrest and impaired cytokine secretion in T-cells (3; **Supplemental Figure 1**).

**Figure 1.**
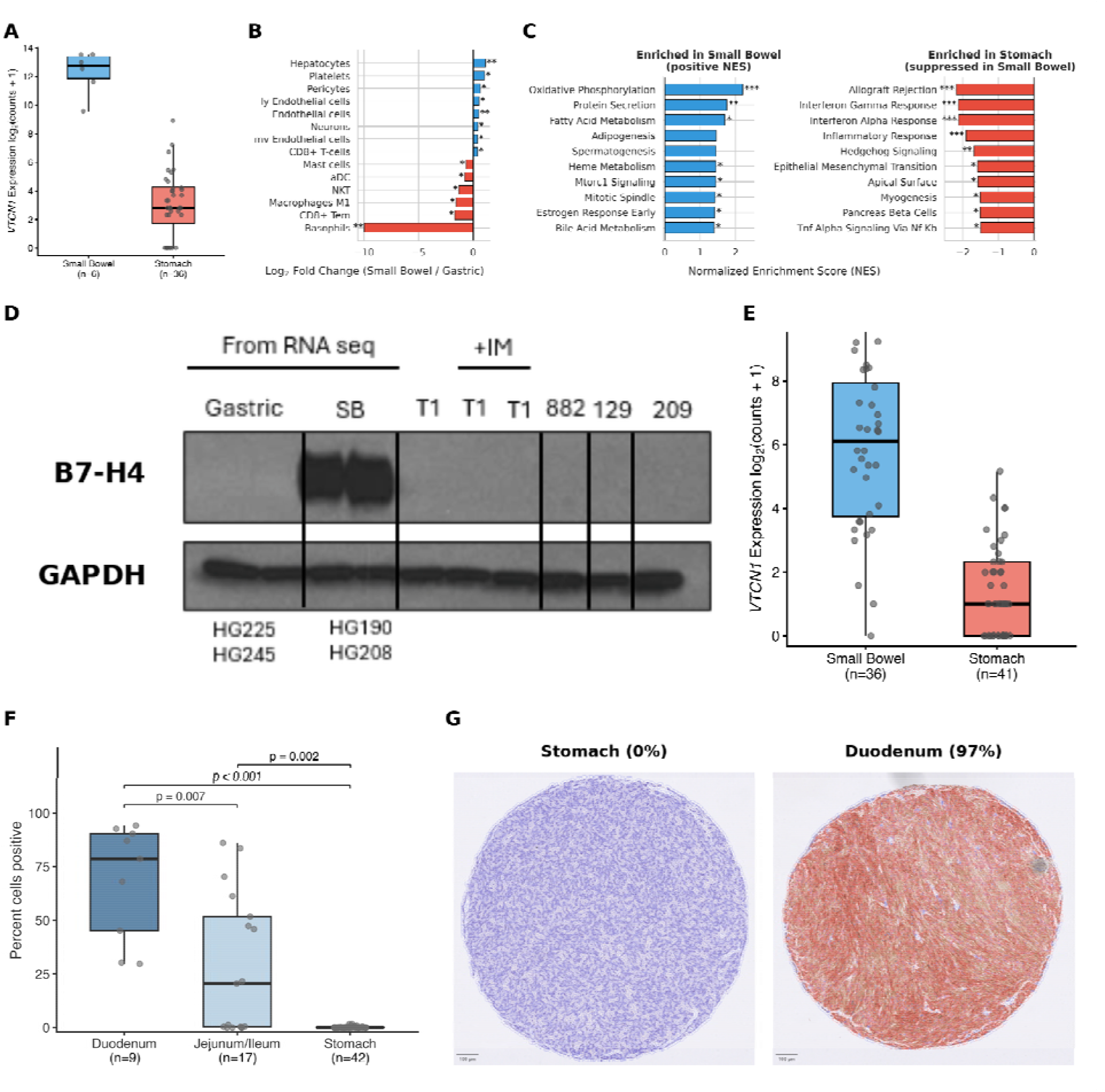
B7-H4 is overexpressed in primary small bowel GISTs and associated with immune suppression. (**A**) *VTCN1* (B7-H4) expression in primary small bowel versus primary gastric GISTs (log_2_FC = 7.95, adj. *P* < 0.001). (**B**) xCell deconvolution of cell-population abundance, small bowel versus gastric GISTs. ly, lymphatic; mv, microvascular; aDC, activated dendritic cells; NKT, natural killer T cells; Tem, effector memory T cells. (**C**) Metabolic enrichment (left; positive normalized enrichment score (NES)) and immune suppression (right; negative NES) in small bowel GISTs. (**D**) Western blot of RNA-seq cohort samples/4 GIST cell lines ± imatinib. SB, small bowel; IM, imatinib; GIST cell lines (T1 and 129, gastric; 882, metastatic; 209, peritoneal metastasis); GAPDH, loading control. (**E**) *VTCN1* expression in small bowel versus gastric GISTs from Gasparotto et al. (5). logLFC = 4.42, *P* < 0.001 by Wilcoxon rank-sum test. (**F**) Percent B7-H4–positive cells in a TMA of 68 untreated primary human GISTs by site, shown as the per-patient mean across 1-3 cores. Median positivity: duodenum 78.6%, jejunum/ileum 20.5%, stomach 0%. (**G**) Representative B7-H4 immunohistochemistry with QuPath classification of a stomach (0%) versus duodenal (97%) GIST core. Red, B7-H4–positive cell; blue, negative cell. Scale bars, 100 µm. In A, E, and F, box bounds show the 25th–75th percentiles, center lines the median, and whiskers the most extreme values within 1.5 × IQR, with all values represented by dots. *<0.05, **<0.01, ***<0.001, denoting nominal *P* (B) or false discovery rate (C).

Gene set enrichment analysis showed suppressed immune signaling in small bowel relative to gastric GISTs. Allograft rejection, interferon-γ and -α responses, and inflammatory response were the most downregulated pathways (all normalized enrichment scores (NES) < −1.9, false discovery rate (FDR) < 0.001), while oxidative phosphorylation (NES = 2.20, FDR < 0.001) and fatty acid metabolism (NES = 1.70, FDR = 0.013) were enriched (**Figure 1C**). The reduced glycolytic dependence may render small bowel GISTs less sensitive to imatinib, which we have shown suppresses glycolysis (4). B7-H4 expression did not differ between imatinib-sensitive and imatinib-resistant tumors (*P* = 0.94), suggesting immune evasion is independent of imatinib resistance and may represent an orthogonal therapeutic target.

Among upstream regulators, *STAT4* and *STAT5B* were overexpressed in small bowel GISTs (log_2_FC = 3.08 and 0.85; adjusted *P* < 0.001 each; **Supplemental Figure 2**). Both correlated with B7-H4 in small bowel tumors (ρ = 0.77 each), but these associations attenuated after controlling for location (*STAT4*: partial *r* = −0.03, *P* = 0.84; *STAT5B*: partial *r* = 0.18, *P* = 0.25), suggesting coordinated site-specific upregulation rather than direct regulation. Since imatinib suppresses STAT signaling, we repeated our analysis on treatment-naïve primary tumors (n=24 gastric, n=5 small bowel). B7-H4, *STAT4*, and *STAT5B* upregulation in the small bowel remained significant (B7-H4: log_2_FC = 9.66, *P* = 0.0006; *STAT4*: log_2_FC = 4.33, *P* = 0.001; *STAT5B*: log_2_FC = 0.93, *P* < 0.001), confirming that site-specific expression differences are independent of imatinib.

Western blot confirmed elevated B7-H4 protein in small bowel versus gastric GISTs in our RNA-seq cohort (n=4). B7-H4 was undetectable in four non–small bowel GIST cell lines regardless of imatinib (**Figure 1D**). We next analyzed bulk RNA-seq data of 77 untreated gastric and intestinal GISTs from Gasparotto et al. (5). *VTCN1* was enriched in intestinal tumors (**Figure 1E**). Immunohistochemistry of our tissue microarray (TMA) of 68 untreated primary GISTs showed median B7-H4 positivity of 78.6% in duodenum, 20.5% in jejunum/ileum, and 0% in stomach (**Figure 1F, 1G, Supplemental Figure 3**). On QuPath classification, B7-H4 staining was diffuse and localized predominantly to tumor cells rather than stroma (**Figure 1G, Supplemental Figure 3**), indicating B7-H4 is likely tumor cell-intrinsic. B7-H4 expression was minimal by immunohistochemistry of 50 gastric GISTs (6).

To our knowledge, we are the first to identify B7-H4 as a site-specific feature of small bowel GISTs, enriched across three independent datasets. This enrichment is associated with an immunosuppressive microenvironment marked by antigen-presenting cell depletion, effector T-cell alterations, and M2-skewed macrophage polarization. These findings may explain why gastric and small bowel GISTs behave differently and support B7-H4-directed immunotherapy for small bowel tumors, with relevance beyond GIST to checkpoint-refractory malignancies. Larger cohorts and functional studies are needed to confirm these associations.

