## Supplemental Figures 1-3, Supplemental Table 1, Methods, for "B7-H4 enrichment represents a site-specific immunotherapy target in small bowel gastrointestinal stromal tumors"

Supplemental Figure 1

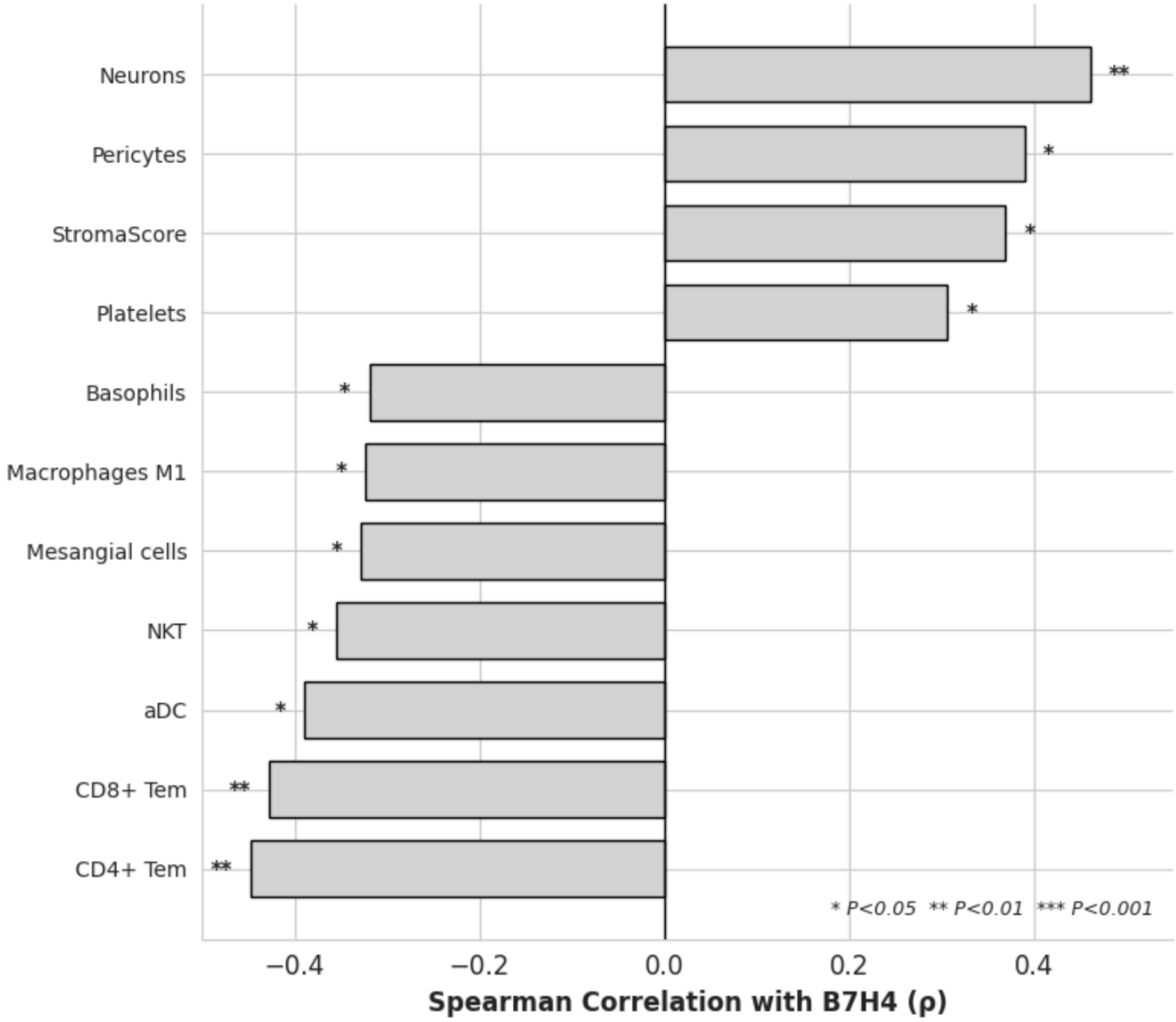


**Supplemental Figure 1.** Spearman correlations between B7-H4 (*VTCN1*) expression and xCell-estimated cell type abundance scores in primary GISTs (n=42). Positive correlations indicate cell types whose abundance increases with B7-H4 expression; negative correlations indicate cell types whose abundance decreases. StromaScore, xCell composite stromal signature score; NKT, natural killer T cells; aDC, activated dendritic cells; Tem, effector memory T cells. *$P$<0.05, **$P$<0.01, ***$P$<0.001.

Supplemental Figure 2

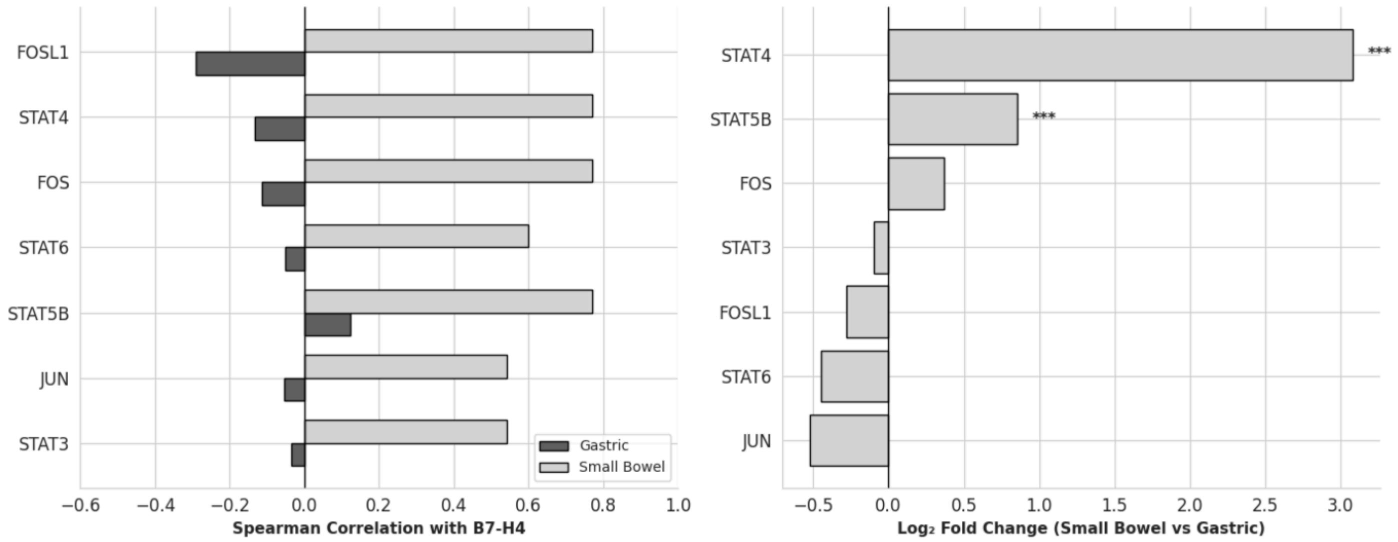


**Supplemental Figure 2.** Location-stratified Spearman correlations between transcription factor expression and B7-H4 (*VTCN1*) in gastric (n=36, dark grey) versus small bowel (n=6, light grey) GISTs (left). Transcription factor differential expression (DESeq2) in small bowel versus gastric primary GISTs (right). $Log_2$ fold-change shown; positive values indicate higher expression in the small bowel. **P*<0.05, ***P*<0.01, *** *P* <0.001.

Supplemental Figure 3

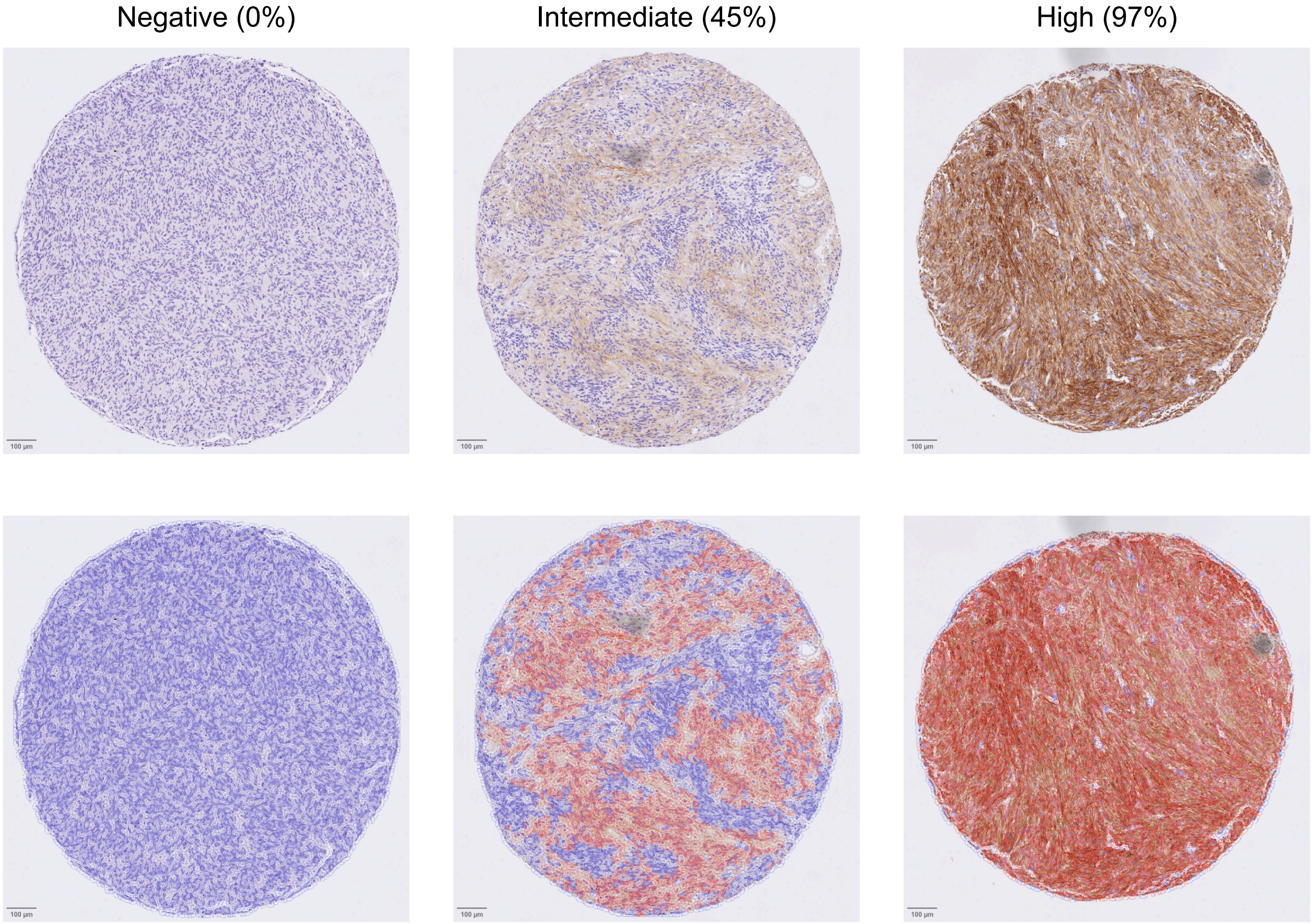


**Supplemental Figure 3.** Representative B7-H4 immunohistochemistry and QuPath cell classification across the dynamic range of staining. Representative examples of tissue core staining in tissue microarray, with range of percent positivity. The 0% is a gastric GIST, the 45% a jejunal/ileal GIST, and the 97% a duodenal GIST. Red annotation indicates a positive cell, and blue a negative cell. Scale bars in bottom left indicate 100 μm.

Supplemental Table 1

| Characteristic | Gastric (n=36) | Small Bowel (n=6) | Total (n=42) |
|---|---|---|---|
| **Treatment status** | | | |
| Untreated | 24 (67%) | 5 (83%) | 29 (69%) |
| Imatinib-sensitive | 6 (17%) | 0 (0%) | 6 (14%) |
| Imatinib-resistant | 6 (17%) | 1 (17%) | 7 (17%) |
| **Mutation status** | | | |
| KIT exon 11 | 16 (44%) | 4 (67%) | 20 (48%) |
| PDGFRA | 13 (36%) | 0 (0%) | 13 (31%) |
| SDH-deficient | 5 (14%) | 0 (0%) | 5 (12%) |
| KIT exon 9 | 0 (0%) | 2 (33%) | 2 (5%) |
| Wild-type | 1 (3%) | 0 (0%) | 1 (2%) |
| KIT exon 13 | 1 (3%) | 0 (0%) | 1 (2%) |

**Supplemental Table 1.** Clinical and pathological characteristics of 42 primary GISTs stratified by anatomical location. Mutation status was determined by targeted sequencing. Values shown as n (%).

### Methods

*Sex as a biological variable*

Sex was not considered as a biological variable.

*Histochemistry*

B7-H4 protein expression was assessed using an antibody clone (clone D1M8I, Cell Signaling Technology). A tissue microarray of 68 untreated primary human GISTs was DAB stained for B7-H4 (rabbit monoclonal D1M8I, Cell Signaling Technology; 1:100) by the University of Pennsylvania Pathology Clinical Service. QuPath was used to quantify percent positive cells per core at an empirically chosen threshold, then averaged across 3 cores (n=57), 2 cores (n=9), or 1 core (n=2) per patient.

*Statistics*

Differential gene expression was analyzed with DESeq2 using default parameters. *P* values were adjusted by the Benjamini-Hochberg method, with significance set at adjusted $P < 0.05$. For gene set enrichment analysis, genes were ranked by the DESeq2 Wald statistic and analyzed with GSEApy. Pathway significance was defined as FDR < 0.25 per GSEA guidelines. Cell type deconvolution was performed with xCell on CPM-normalized data, which scores 64 cell types and 3 composite signatures. Cell type abundances were compared between anatomic sites with Mann-Whitney U tests and are reported as nominal *P* values, without correction for multiple comparisons. Transcription factor differential expression was also assessed with DESeq2 and Benjamini-Hochberg correction. Correlations with B7-H4 expression were assessed with Spearman's rank correlation. Partial correlations controlling for anatomic location were computed as the Pearson correlation of residuals after each variable was regressed on site. Location-stratified correlations in the small bowel (n = 6) are exploratory given the small sample. B7-H4 expression and imatinib response were compared with the Mann-Whitney U test. In the tissue microarray, percent B7-H4–positive cells were compared across sites with pairwise Wilcoxon rank-sum tests, without correction. *VTCN1* expression in the Gasparotto et al. cohort was compared with the Wilcoxon rank-sum test, which is equivalent to Mann-Whitney U. Unless otherwise noted, *$P < 0.05$, **$P < 0.01$, ***$P < 0.001$. Analyses were performed in Python and R.

*Study Approval*

Tumor specimens were obtained from patients who provided written informed consent prior to surgery, in accordance with a protocol approved by the University of Pennsylvania Institutional Review Board (IRB #852459).

*Data Availability*

Raw RNA sequencing data and metadata from Vitiello et al. *J Clin Invest* 2019 are publicly available for download from the NCBI Sequencing Read Archive (SRA) under accession no. PRJNA521803. Raw RNA-sequencing data from Gasparotto et al. *JCI Insight* 2020 are accessible at the NCBI-SRA database (accession PRJNA637476). Values for all data points in graphs are reported in the Supporting Data Values file.

*Acknowledgments*

We would like to acknowledge Gasparotto et al. for providing the metadata from their cohort of GISTs, which helped validate our findings of enriched B7-H4 expression in small bowel GISTs and confirm generalizability for research in the larger GIST community.
